# Trait-specific chromatin architectures channel pleiotropic genes toward sexually dimorphic development in horned beetles

**DOI:** 10.64898/2026.03.19.712098

**Authors:** Erica M. Nadolski, Armin P. Moczek

## Abstract

Sex-responsive trait development generates much of the phenotypic variation found in natural populations and diversifies rapidly among closely-related taxa. Furthermore, rather than exhibiting equal sexual dimorphism across all traits, organisms are mosaics of tissues that vary in their degree of dimorphism. Yet, how these mosaic patterns are generated remains largely an open question, as sexually dimorphic traits have typically been studied individually in select model systems. In this study, we compare gene regulatory landscapes across five traits that differ in the degree of morphological sexual dimorphism in the bull-headed dung beetle *Onthophagus taurus* by assaying tissue-specific gene expression and chromatin accessibility at the onset of pupal development when future adult form is specified. We identify a modest number of pleiotropic regulators associated with sex differences across traits, yet uncover a high degree of sex- and trait-specificity in chromatin architecture within developing tissues. We then confirm the role of the sex determination factor *doublesex* in the regulation of sex differences through expression of sex-specific isoforms, and uncover trait- and sex-specific sets of Doublesex binding sites likely underpinning context specific sexual dimorphisms. Further, we identify and functionally validate the transcription factor ventral veinless as a regulator of sexually dimorphic development. Our findings suggest that in contrast to *doublesex, ventral veinless* does not exhibit sex-biased expression, yet exerts its sex-specific regulation via sets of differentially accessible binding sites. This work furthers our understanding of the molecular mechanisms instructing the development of sex differences and provides novel insights illustrating how transcriptional activity and chromatin remodeling interact to generate sexual dimorphism in a trait-specific manner. More generally, our work contributes to a growing body of knowledge on how development integrates cues such as sex determination to enable highly similar genomes to yield diverse phenotypic outcomes.

## Introduction

Sexual dimorphism, differences between males and females of a species, is a major axis of intraspecific variation and diversifies rapidly among taxa across the tree of life. Sexual dimorphism is thought to reflect adaptive divergence in response to selection favoring different optimal character states in males and females (Andersson 1994). However, other than the few genes located on the sex-limited copy of heteromorphic sex chromosomes (such as the Y chromosome in XY systems), there are no genes that exist exclusively in one sex. Instead, sex-biased expression of the same genes emerges as the primary mechanism to resolve conflicts between sex-specific adaptive peaks, allowing each sex to approach its own adaptive optimum independently (Ellegren and Parsch 2007, Williams and Carroll 2009). While the ecological and fitness relevance of sex-dependent development is often well characterized, the genetic and developmental mechanisms underlying sexual dimorphism and its evolution are not. Even though sexual dimorphisms have been studied from diverse perspectives and key regulators of sex-specific development have been identified, such as *doublesex* in insects (Shukla and Nagaraju 2010) and its ortholog DMRT in mammals (Kopp 2012), our understanding of how sexually dimorphic trait expression is instructed throughout development and diversifies across traits and species remains largely incomplete.

Additionally, rather than exhibiting equal sexual dimorphism across all traits, organisms are mosaics of tissues that vary in their degree of dimorphism (Andersson 1994). Even though sex differences across different body regions are likely to be shaped at least in part by unique selection pressures, there remains a finite number of pre-existing gene regulatory networks available to respond to these selection pressures and scaffold the development of sex differences across the body. Thus, shared molecular underpinnings of sexual dimorphism may be a key nexus whereby development may constrain, bias, or otherwise shape responses to sex-specific selection pressures. Yet, outside of a few key regulatory genes such as the insect sex determination factor *doublesex* that have been systematically studied across tissues, timepoints, and taxa (Burtis and Baker 1989, Kopp 2012, Clough et al. 2014), the degree to which sex-biased gene expression and regulation are shared across developmental contexts remains largely an open question. Some theoretical predictions from sexual selection theory suggest that sex-biased gene expression should be shared across traits to enable coordinated transcriptomic responses to a global cue such as sex (Cheverud 1996). However, alternative predictions have been made that sex-biased gene expression should instead generally be trait-specific in order to alleviate negative pleiotropic constraints (Hodgkin 2002).

The transcriptional basis of sexual dimorphism remains poorly understood outside select model systems and traits. Further, it is well known that the interplay between transcription factor activity and nucleosome occupancy at *cis-*regulatory regions is key to establishing specific regulatory states (Loker et al 2021, Davidson et al 2022), yet the nature of this process remains to be fully integrated into our understanding of the regulation of sex differences. Here we use tissue-specific RNA-seq and ATAC-seq to characterize gene expression and chromatin accessibility in the bull-headed dung beetle, *Onthophagus taurus*, a representative of a clade that possesses striking and varied morphological sex differences across multiple traits. Specifically, we compare sex-responsive gene expression and chromatin accessibility in the pupal tissue primordia of five traits that differ in the degree of sexual dimorphism (Figure 1A): (i) external components of **genitalia** develop from homologous larval tissues into strikingly different adult traits (ii) **posterior head horns** are phylogenetically widespread, secondary sexual weapons employed in male-male mate competition. The ancestral state for the clade includes horned males and hornless females, but male horn size has been greatly exaggerated in *O. taurus* (Emlen et al. 2005). (iii) **The anterior head** is also used in fighting maneuvers between males during competition. Female *O. taurus* possess a prominent horizontal ridge across the clypeus, mirroring the most common phenotype in their clade, while males have no such ridge but instead possess an upturned lip at the front of the clypeus. (iv) **Fore tibiae** are also sexually dimorphic to a mild degree, reflecting alternative behavioral adaptations: forelegs are used primarily for tunnel digging by females, who possess wider limbs with more prominent teeth, while males also use their fore tibiae in courtship to drum on female elytra, and possess narrower limbs with thinner teeth (Rohner et al. 2023). Finally, (v) **fore wings** represent a monomorphic trait in this species and exhibit no obvious sex differences in adult beetles. Thus, the array of sexually dimorphic traits in *O. taurus* exist on a spectrum of varying *magnitude* of sex differences. In addition, the unique evolutionary history of each trait offers a second spectrum on which the gene regulatory basis of sex-biased development can be compared: for example, genitalia represent a sexually dimorphic trait class in existence for hundreds of millions of years, predating the origin of beetles, including the genus *Onthophagus*. In contrast, sex differences in the posterior head, anterior head, and fore tibia are much more recently evolved and restricted to small clades of dung beetle species. In fact, the posterior head horns of males fit even the strictest definition of an evolutionary novelty, exhibiting no homology to existing traits in ancestral lineages (Linz and Moczek 2020). Thus, we hypothesized that the rapid evolution and diversification of novel and exaggerated sex differences in *O. taurus* may be facilitated by the reuse and recombination of pre-existing gene regulatory mechanisms operational in the development of other sexually dimorphic traits.

**Figure 1.**
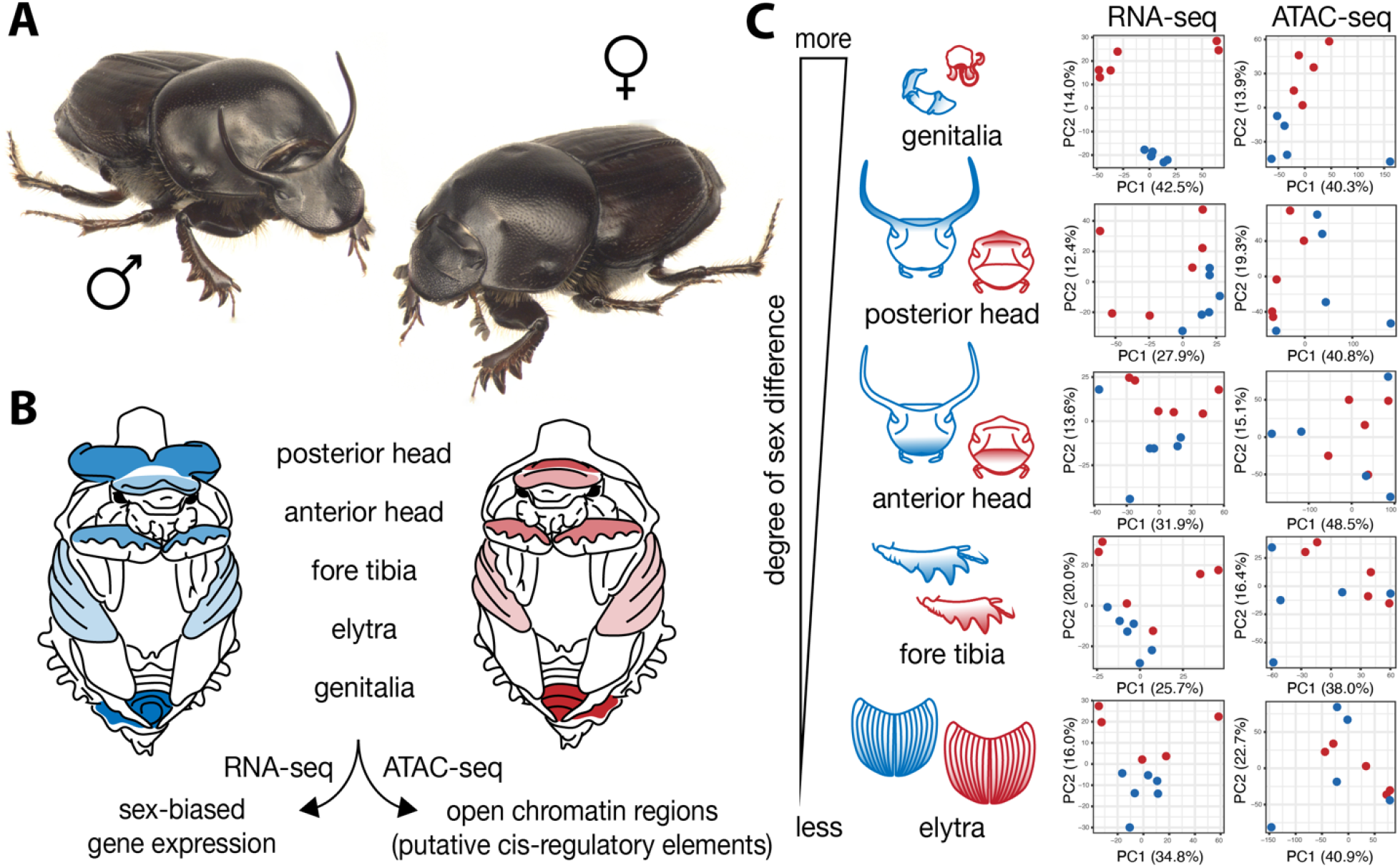
Sex and tissue type drive global patterns of gene expression and chromatin accessibility in sexually dimorphic adult *Onthophagus taurus* beetles. (A) Adult male (left) and female (right) *O. taurus*. (B) DNA and RNA were extracted from the epithelial tissue of the posterior head, anterior head, fore tibia, elytra, and external components of the genitalia from animals within 12 hours of pupation, depicted in a male (blue) and female pupa (red). (C) Principal component analysis plots showing the distribution of variation along PC1 and PC2 across biological replicates for each body region after RNA sequencing and ATAC sequencing, with male samples shown in blue and female samples shown in red. The relative degree of morphological difference between the sexes for each trait is indicated by the gradient on the right.

Here, we assayed sex-responsiveness in gene expression and chromatin accessibility during the development of traits spanning a wide range of degrees of sexual dimorphism to unveil the molecular basis of how development generates sexually dimorphic phenotypes, and how these processes are employed to adjust development across different body regions. We predicted that the *number* of sex-responsive genes and chromatin regions would scale with the magnitude of morphological sex differences across traits. Through our comparative approach, we assessed whether sexually dimorphic development across the body is enabled by a shared core of pleiotropic genes and pathways or alternatively by primarily trait-specific repertoires of genes and *cis-*regulatory elements. Finally, we employed functional genetics to uncover the co-option of a conserved regulator of epithelial patterning into the regulation of sex differences across multiple traits in *O. taurus* and propose a mechanism by which a transcription factor can affect the sexes uniquely without being differentially expressed. Combined, this work sheds light on the specific and synergistic roles of trait- and sex-biased gene expression and chromatin accessibility in regulating sex-biased development and its diversification among traits. These findings contribute to a broader understanding of how development can integrate cues such as sex determination to enable highly similar genomes to yield varying adaptive morphologies.

## Results and Discussion

Sexual selection is credited with having driven the extraordinary diversity of exaggerated secondary sexual traits such as weapons and ornaments (Andersson 1994, Lavine et al. 2015). Understanding the genetic architecture underlying the development of these traits is of interest to diverse disciplines. For example, sexual dimorphic trait formation affords developmental geneticists a window into mechanisms that enable a single genome to integrate diverse inputs toward context-specific, adaptive developmental outcomes. At the same time, evolutionary and evolutionary developmental biologists can harness the degree and diversity of sexual dimorphisms to understand how these proximate mechanisms may enable, bias and constrain trait diversification. However, this topic has been underexplored outside of select model systems and traits, and most work has been restricted to the level of gene expression differences, leaving unexplored the mechanisms by which differential gene expression arises in the first place, let alone how it evolves as a lineage diversifies. We leveraged the diversity of sexually dimorphic traits in the bull-headed dung beetle *O. taurus* to first test a hypothesis arising out of sexual selection theory which predicts that the sex exhibiting more dramatic trait elaboration should also express more sex-biased genes (Levine and Tjian 2003). We then probe one of the mechanisms proposed to underpin such gene expression differences by testing the hypothesis that different sexually dimorphic traits will exhibit distinct patterns of chromatin accessibility, thereby providing a proof of principle that tissue- and sex-specific chromatin remodeling is a major mechanism facilitating the regulation of sexually dimorphic development across the body. Using genome-wide RNA sequencing (RNA-seq) and ATAC sequencing (ATAC-seq), we simultaneously profiled gene expression and chromatin accessibility in both sexes across five body regions that differ in the degree of sexual dimorphism (Figure 1 A-B). We then used these datasets to assess the degrees of relative pleiotropy in both sex-responsive genes and open chromatin regions (OCRS), distinguishing between shared and trait-specific regulators, OCRs, and developmental mechanisms. Finally, we functionally validate the role of one newly-identified candidate regulator in regulating the development of sex differences across the traits studied here and highlight its noncanonical mode of action.

### Sexually dimorphic growth requires large-scale differential chromatin accessibility and gene expression

We first identified 15,979 expressed transcripts across all trait samples through tissue-specific RNA-sequencing. The global patterns of gene expression variation across samples are driven primarily by trait (Supplementary Figure 1), followed by sex, with the sexes clustering visually when plotted onto PC1 and PC2 for the majority of tissue types (Figure 1C). To probe the level of sex-responsiveness in the transcriptome, we tested for statistically significant differences in gene expression levels between the sexes for each trait. We predicted that the relative degree of sex-responsiveness in the transcriptome should scale with the degree of morphological sex differences when compared across traits (G > PH > AH > FT > E). This prediction was not met; instead the patterns uncovered here showed total numbers of DE genes being marginally highest in the anterior head and lowest in the fore tibia (AH > G > PH > E > L). The high number of sex-responsive genes in all traits was surprising; the total number of DE genes was within an order of magnitude across all traits and independent of the degree of morphological sex differences therein (Figure 2A-B, supplementary figure 2, Supplementary table 1). These findings are also at odds with predictions from sexual selection theory positing that the sex exhibiting more dramatic trait elaboration should also express more upregulated genes, or in other words, that morphological and transcriptional complexity should scale together (Levine and Tjian 2003, West-Eberhard 2003, Carroll 2008, Wyman et al 2012). Instead, our results indicate that the regulation of sex-responsive developmental processes requires large-scale upregulation of genes in both sexes, independent of the magnitude of morphological sex-differences that arise as development culminates, suggesting that active repression of alternative developmental outcomes may be a common molecular phenomenon.

**Figure 2.**
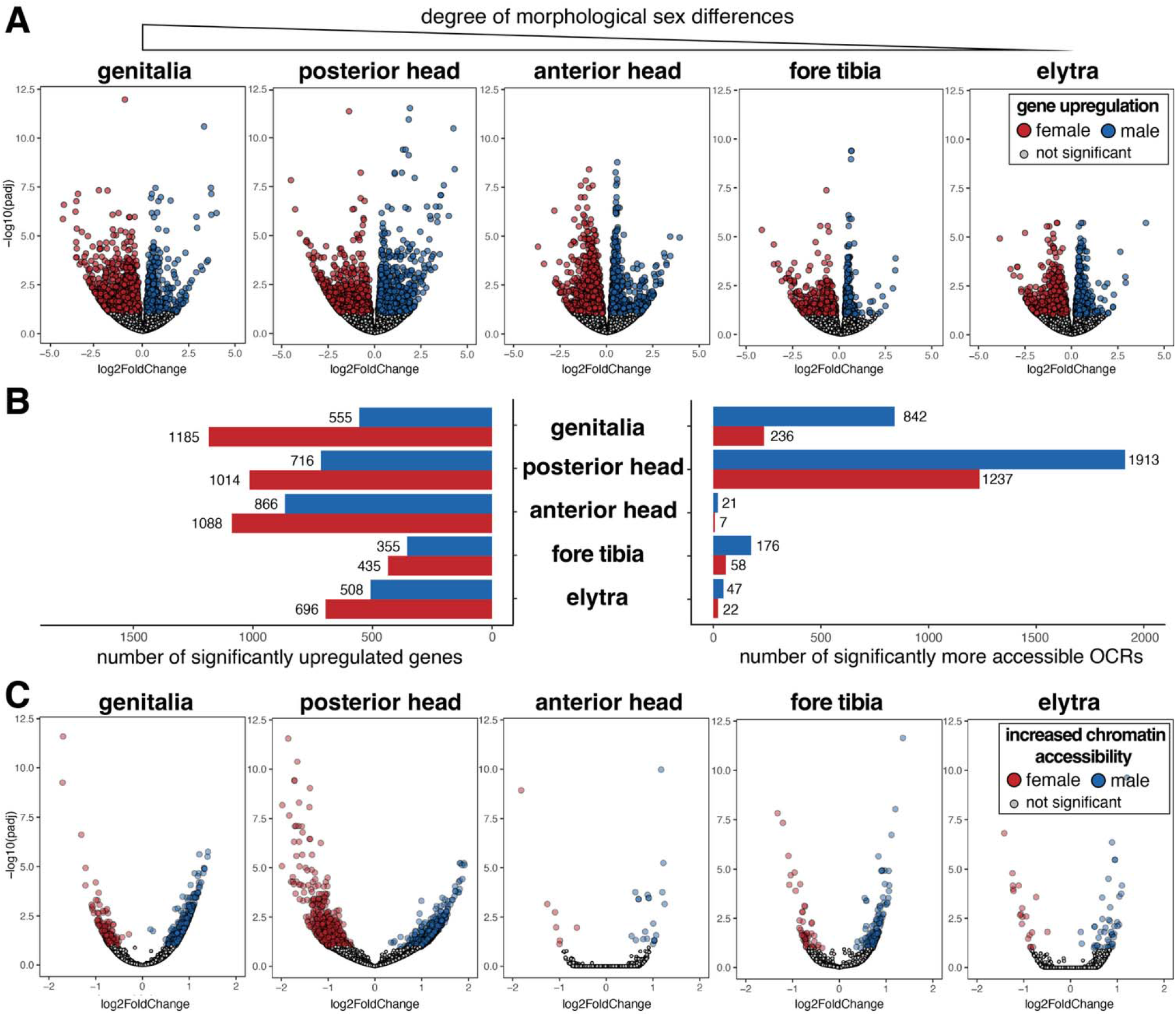
Degree of sex bias in chromatin accessibility, but not in the transcriptome, maps onto the relative degree of morphological sexual dimorphism for a given body region. (A) Volcano plots of expressed genes from each body region, with dots colored in red and blue exhibiting significantly upregulated expression in females or males respectively. (B) Bar charts tallying the total number of differentially expressed genes (left) and differentially accessible OCRs (right) across all body regions. (C) Volcano plots of open chromatin regions from each body region, with dots colored in red or blue exhibiting a significant degree of greater accessibility in females or males, respectively.

Next, we sought to probe one of the mechanisms by which these gene expression differences arise. To do so, we identified 88,405 high-confidence OCRs across all trait samples through tissue-specific ATAC-sequencing at the same developmental stage. Mirroring the transcriptomic patterns, the global patterns of variation in chromatin accessibility across samples were also driven primarily by trait type, followed by sex (Supplementary Figure 1). However, when plotted on PC1 and PC2, the sexes visually cluster only in the genitalia and posterior head comparisons (Figure 1C). We tested for statistically significant differences in chromatin accessibility between the sexes for each body region, again predicting that the relative degree of sex-responsiveness in chromatin accessibility should scale with the degree of morphological sex differences (G > PH > AH > FT > E). The numbers of differentially accessible (DA) OCRs were highest for the posterior head followed by genitalia, the two traits with the most dramatic morphological differences, while the corresponding numbers were an order of magnitude lower for the remaining three much *less* sexually dimorphic traits (Figure 2B-C, Supplementary figure 3, Supplementary table 2). Intriguingly, the posterior head tissue showed a three-fold higher number of DA OCRs than the genitalia, despite the extraordinary degree of sex bias inherent in insect genitalia formation. However, despite similar degrees of morphological dimorphism in both traits, the sexual dimorphism in genitalia is ancient, predating even the origin of beetles, while the sex differences in the posterior head have evolved much more recently in the subfamily Scarabaeinae, within the last 86-100MY (Ahrens et al 2014). Thus, the patterns uncovered here may suggest that the rapid diversification of conditional development – such as the sex-specific morphological exaggeration of male posterior head horns – can be enabled in large part by changes to chromatin architecture.

**Figure 3.**
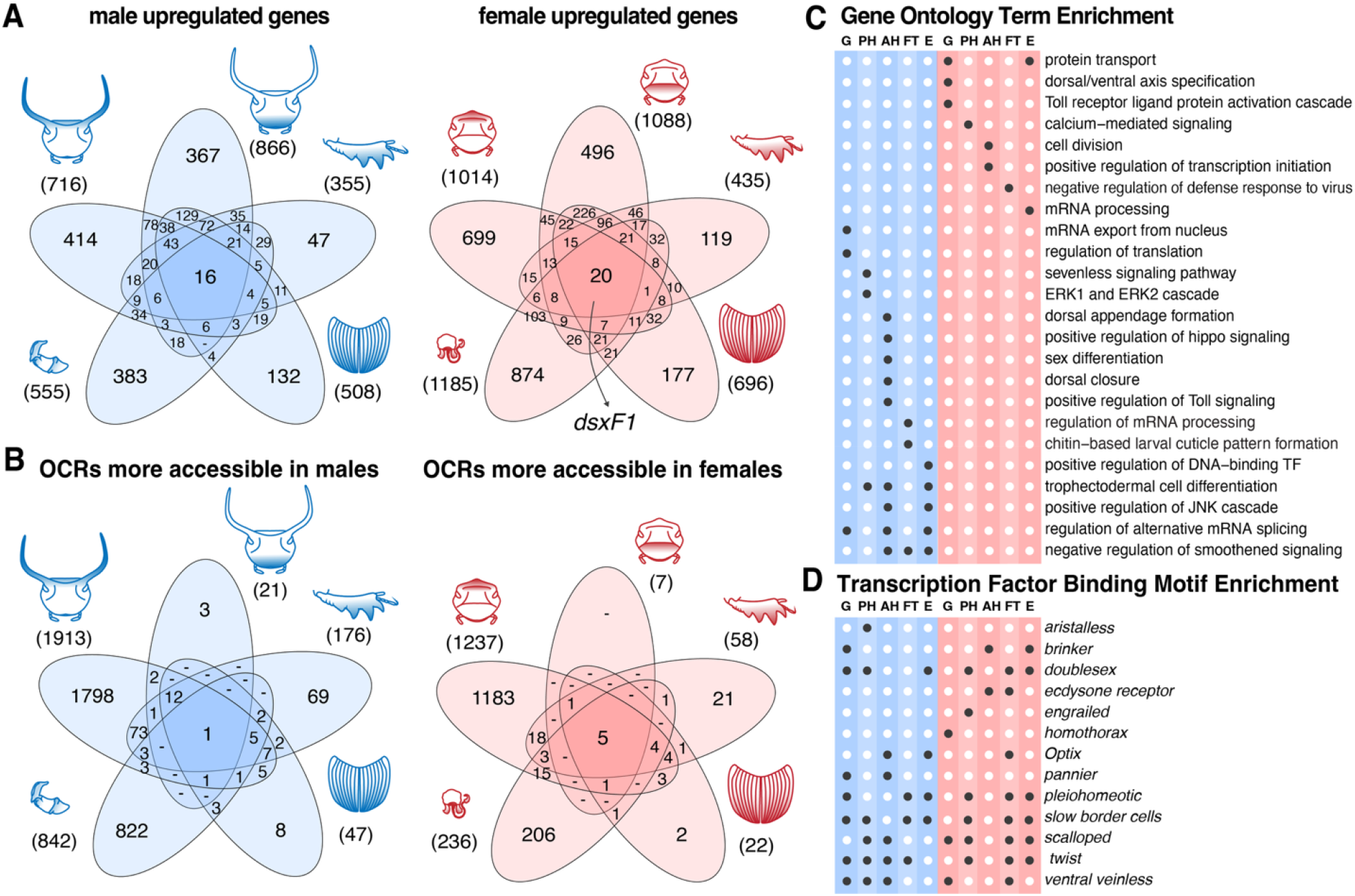
Sex-responsive chromatin accessibility exhibits high levels of trait-specificity, while sex-biased genes and pathways are shared to a much greater degree across developing traits. (A) Venn diagram showing genes that exhibited elevated expression in one or more traits in males (blue) or in females (red). (B) Venn diagram showing open chromatin regions that exhibited greater accessibility in one or more traits in males (blue) or in females (red). (C) Plot of enriched gene ontology terms associated with upregulated gene expression patterns in males or females for a given body region with enrichment of *p*<000.1 indicated by dark circles. (D) Transcription factor binding motif enrichment was analyzed for the sets of sex-responsive OCRs in each trait. Examples of transcription factors showing enriched binding motif frequencies are shown here; enrichment of *p*<000.1 is indicated with dark circles and genes are ordered alphabetically. G = genitalia; PH = posterior head; AH = anterior head; FT = fore tibia; E = elytra.

Taken together, our predictions regarding the molecular patterns underlying morphological sex differences were only supported for a subset of comparisons. In particular, our results suggest that the regulation of sex-responsive developmental processes appears to require large-scale shifts in gene regulation and chromatin accessibility in both sexes, regardless of the degree of morphological trait exaggeration occurring in each. While at odds with predictions from sexual selection theory, previous studies of sex-biased gene expression have reported a similar pattern in the same species as well as other insect taxa (Ledon-Rettig and Moczek 2016, Zinna et al. 2018). Notably, across all traits assayed, females upregulated a greater number of genes than males, while males exhibited a greater degree of chromatin accessibility than females. Lastly, our findings suggest that even the formation of traits that appear monomorphic on a morphological level may be underlain by cryptic sex-biased physiological and developmental processes, as analysis of sexually monomorphic elytra samples revealed numbers of sex-responsive genes and OCRs similar to foretibiae and anterior heads, traits exhibiting obvious sex-specific morphologies.

### Pleiotropy in gene expression, trait-specificity in OCR accessibility

Next, we sought to assess the degree to which the sets of sex-responsive genes and OCRs scaffolding the development of sexual dimorphism might be *shared across traits*. Some theoretical work (e.g. Cheverud 1996) predicts that sex-biased gene expression should be shared across traits to enable coordinated transcriptomic responses to global cues such as sex, while competing theories (e.g. Hodgkin 2002) have posited that sex-biased gene expression should instead be generally trait-specific in order to alleviate negative pleiotropic constraints. Here we sought to address these contrasting perspectives at two levels of molecular gene regulation-differential gene expression and differential chromatic accessibility. To our knowledge, these theoretical predictions have not previously been tested using chromatin accessibility data. Further, we sought to test whether patterns of trait-specificity would emerge more strongly on the level of chromatin accessibility than on the level of gene expression, as studies of gene regulation have established that modularity in gene regulatory networks is often achieved by the evolution of temporally- and spatially-specific enhancers (Gompel et al. 2005, Williams et al. 2008, Rice et al. 2019).

For these analyses, we focused on the sets of upregulated genes and differentially accessible OCRs *within each sex* across tissues. First, we assessed the degree of overlap in the sets of male- and female-biased genes detected across each of our five focal traits (Figure 3A). Of the 3000 total male-biased genes, 1343 (44.8%) were upregulated in only a single trait, and of the 4418 female-biased genes, 2365 (53.5%) were upregulated in one trait only. Across both sexes, nearly half of all sex-responsive genes were upregulated in two or more traits, indicating a high degree of shared molecular underpinnings driving sex-biased development across distinct body regions. Furthermore, 16 male-biased genes and 20 female-biased genes were significantly upregulated across all five traits, suggesting that genetic pleiotropy is not uncommon for regulators associated with sex-biased development, a finding that has also been reported in previous transcriptomic studies of insect sexual dimorphism (Ledon-Rettig and Moczek 2016, Zinna et al 2018). In contrast, and in support of our predictions, the degree of overlap in sex-responsive chromatin architecture across traits was much lower than on the level of gene expression: of 2999 total male-biased OCRs, 2700 (90.0%) were sex-responsive within one trait, and of 1560 female-biased OCRs, 1412 (90.5%) were unique to one trait only (Figure 3B). The high degree of trait- and sex-specificity seen on the level of chromatin architecture suggests a plausible mechanism for channeling the functions of otherwise pleiotropic regulators into context-specific developmental outcomes.

Next, we sought to extend these investigations into shared developmental processes through gene ontology (GO) enrichment analyses on the sets of male- and female-upregulated genes from each trait. On average, enriched GO terms from the sets of genes upregulated in males outnumbered female GO terms roughly 2:1 (Supplementary table 3). While many GO terms were enriched in only single sets of sex-responsive genes, the terms that were enriched across more than one gene set offer some insight into the shared processes that may be underlying sex-biased development across the body in *O. taurus* (Fig 3C). Male-biased GO terms enriched in more than one trait included: regulation of alternative mRNA splicing, trophectodermal cell differentiation, negative regulation of smoothened signaling, and positive regulation of JNK cascade. These shared processes suggest that conserved signaling pathways may have been co-opted into the regulation of sex-biased development across multiple body regions, either by recruitment of genes and pathways whose expression was already ancestrally sex-biased, or convergent selection upon the same pathways *de novo* across each trait, or a combination of the two.

In general, the analyses above suggest that sex- and trait-responsive chromatin architecture may be a key mechanism for channeling pleiotropic gene expression toward context-specific sex-biased developmental outcomes. Thus, we sought to investigate what pleiotropic regulators might be acting as key nodes within such gene regulatory networks. To do so, we performed transcription factor binding motif enrichment analyses using the sets of sex- and trait-responsive OCRs identified earlier. Motif enrichment analyses identify binding sites that show significantly increased frequencies within a target set of genomic regions (here being the set of male- or female-biased OCRs from one trait) relative to the sites’ frequencies in a background set of genomic regions (here being the total set of OCRs across all trait types). These analyses implicated a number of transcription factors – some of which were enriched in just one developmental context, such as *aristaless, engrailed*, and *homothorax* – while others were implicated across a variety of contexts, such as *doublesex, scalloped*, and *twist* (Figure 3D, supplementary table 4). Importantly, these results identify candidate regulators that may be binding context-specific OCRs, and thus any such regulators showing enriched binding motif frequencies across multiple sets of sex-responsive chromatin may represent particularly interesting candidates for future functional studies.

### Trait- and sex-responsive chromatin architectures channel the functions of the insect sex determination factor *doublesex*

The transcription factor *doublesex* (*dsx*) is conserved across insects as the downstream effector of the sex determination pathway (Shukla and Nagaraju 2010, Kopp 2012, Bopp 2013, Mitchell et al. 2025), and has also been implicated as a developmental regulator of sex-biased phenotypes across diverse insects (Clough et al. 2014, Deshmukh et al. 2020). Because insects undergo cell-autonomous sex determination, it was traditionally thought that *dsx* must be expressed across all cell types, but recent work has shown *dsx* expression to instead be highly context-specific (Verhulst and van de Zande 2015). Thus, its widespread involvement in the regulation of sex-biased traits – particularly in those that can be considered evolutionary novelties – suggests that *dsx* must have been repeatedly co-opted into new developmental contexts. The general conservation of *dsx* function in onthophagine beetles has previously been established; *dsx* gene expression knockdown eliminates somatic sex differences across the body by generating males and females sharing intermediate phenotypes, including the four sexually dimorphic traits studied here (Kijimoto et al. 2012, Rohner et al. 2021). However, it remains unclear how *dsx* exerts its regulatory functions in a trait-specific manner, or how it has been repeatedly co-opted into new body regions as sex differences first arose.

Given the key role of *dsx* as a sex determination and differentiation factor in insects, we sought to characterize the chromatin architectures surrounding both the *dsx* locus and the loci of its putative target genes. As has been shown across diverse insect species, transcripts from the *dsx* locus in *O. taurus* are spliced into sex-specific isoforms (Kijimoto et al. 2012) which regulate partially overlapping sets of target genes (Ledon-Rettig et al. 2017). The short-read transcriptomic data generated here uncovered statistically significant upregulation of the longest female isoform *dsxF1* across all traits in female tissues as expected, although the relative expression level of the isoform differed across traits (Fig 4A), consistent with the necessity of this gene in establishing sexual identity on an intracellular level before sex differentiation can occur. At the same time, our transcription factor binding site enrichment analyses also uncovered significant enrichment of putative Dsx binding motifs across a range of sex- and trait- responsive chromatin regions (Fig 4B).

**Figure 4.**
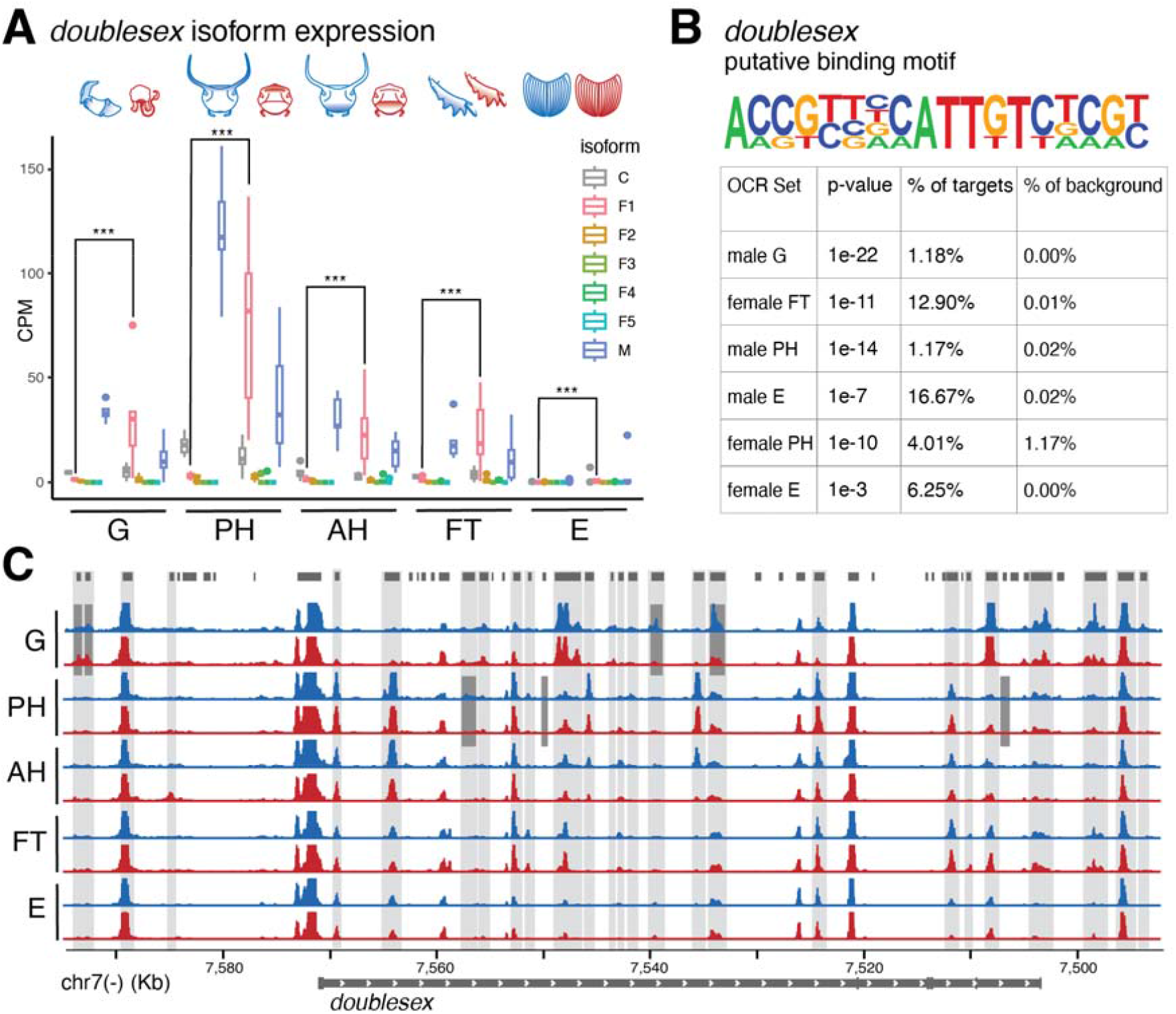
An established regulator of sexual dimorphism shows trait- and sex-biased isoform expression and chromatin architecture. (A) The transcription factor Doublesex is spliced into sex-specific isoforms, some of which show consistent differential expression across all traits assayed. C = common isoform; F1-F5= female isoforms 1-5; M = male isoform. (B) The binding motif associated with the DNA-binding domain shared across Doublesex isoforms is enriched in the sets of sex-responsive chromatin regions from multiple traits. (C) Many open chromatin regions around the *doublesex* locus (top track) show patterns of differential accessibility across traits (light grey bars) and between the sexes (dark gray bars). G = genitalia; PH = posterior head; AH = anterior head; FT = fore tibia; E = elytra.

In combination, these findings indicate that regulatory sex differences exist at two distinct levels surrounding the Dsx ‘node’ of the sex differentiation network in horned beetles: sex differences in the *trans* regulator Dsx itself generated by sex-specific splicing, in addition to sex differences in the *cis*-regulatory elements accessible for the protein to bind. Lastly, we assessed the chromatin architecture within 25kb up and downstream of the *dsx* locus and found 7 sex-responsive elements and 51 trait-responsive elements, suggesting that differential cis-regulatory element accessibility may be one mechanism by which *dsx* transcription is regulated in a context-specific manner during development (Fig 4C, Supp data table 6). Indeed, transgenic work in flies has mapped separate, modular intronic enhancers responsible for driving *dsx* expression in the sexually dimorphic organs of the fly foreleg (Rice et al. 2019). The existence of trait-specific, combinatorial cis-regulatory elements regulating *dsx* expression could also explain the independent co-option of *dsx* regulation into new body regions. Future work investigating the potential upstream regulators binding these putative trait-specific regulatory elements at the *dsx* locus may be able to provide insight into how this gene has been repeatedly co-opted into the regulation of sexual dimorphic development in new body regions, and whether this has occurred as part of larger gene regulatory network co-option or the evolution of *de novo* regulatory interactions.

### Co-option of *ventral veinless* as a pleiotropic regulator of sexually dimorphic development

The analyses described above highlighted a number of genes as potential regulators of sexual dimorphism. We were particularly interested in genes encoding transcription factors, and even more so in the subset that emerged with significant associations to sex differences in more than one trait. An increasing number of studies indicate that co-option of conserved regulatory genes into new ontogenetic contexts is one of the major mechanisms underlying the origin and diversification of form (True and Carroll 2002, Martin et al. 2014, Glassford et al. 2015). Many of the candidate regulators identified though this study showed patterns of sex-biased expression *in combination with* patterns of sex-biased binding site accessibility in one or more body regions, e.g. *doublesex* (Figure 4). However, several additional candidates showed patterns of sex-biased binding site accessibility *without* being differentially expressed themselves, including *scalloped* and *ventral veinless* (Figure 6A). We were intrigued by this class of regulators and sought to assess whether a transcription factor displaying such genomic patterns could be playing a role in regulating the development of sex differences.

To formally test the role of the candidate regulator *ventral veinless (vvl)* in regulating sexually dimorphic development in *O. taurus*, we experimentally reduced *vvl* expression levels using RNA interference (supplementary tables 5, 6). Previous work in fruit flies has implicated *vvl* in the regulation of wing size and venation (de Celis et al. 1995), in addition to roles in embryonic nervous system patterning and adult immunity (Meier et al. 2006, Junell et al. 2010). *Ot-vvl*^RNAi^ treatment at a range of dosage concentrations resulted in adult beetles with diverse morphological differences compared to controls. Fore tibiae were affected similarly in both males and females: overall length decreased in all RNAi beetles, as did the spacing of tibial teeth; in the most strongly affected individuals tibial teeth were largely absent (Figure 6B). In the anterior head in contrast, *Ot-vvl*^RNAi^ yielded unique, sex-specific phenotypes, in a manner that appeared to reverse wild-type sex-biased morphology. Specifically, the anterior ridge seen only in control females was deleted in RNAi females and was replaced with an upturned clypeal lip normally seen only in control males, while the clypeal lip disappeared in RNAi males and was instead replaced by a subtle ridge resembling wild-type and control female morphology (Figure 6C; note that the body color and male horn size variation present in this figure represents naturally-occurring variation due to body size and does not reflect phenotypes resulting from RNAi treatment). In addition, the typically smooth shape of the wild-type male pronotum was transformed toward a morphology resembling females, with RNAi males exhibiting small bilateral protrusions (Figure 6C). Although previous work suggests that wing patterning should be affected, we were unable to systematically assess wing and elytra size and venation in *Ot-vvl*^RNAi^ adults, as treatment prevented the successful molting and shedding of the pupal cuticle from the entire body in the majority of individuals, likely leading to morphological defects in wing morphogenesis due to physical constraints imposed by incomplete ecdysis alone. These results functionally implicate the transcription factor *vvl* in the regulation of sexually dimorphic development in horned beetles, and suggest a model wherein sex-specific development is instructed by a gene regulatory network not through gene expression level differences but via differential accessibility of key binding sites (Fig 6D). Furthermore, these results suggest that the ancestral epithelial patterning function of *vvl* in the context of wing development was secondarily coopted into the regulation of multiple novel sex differences in beetles.

## Conclusions

Sex-responsive trait development generates a major fraction of the phenotypic variation found in natural populations, and the resulting sexually dimorphic traits rank among the most rapidly evolving trait categories. Yet how sexual dimorphism initially emerges from the existing regulatory architecture scaffolding monomorphic trait development remains largely unresolved. In this study, we compare gene regulatory landscapes across five traits that differ in the degree of morphological sexual dimorphism in the bull-headed dung beetle (Fig. 2). We identify a small number of pleiotropic regulators of sex differences across these traits, and at the same time uncover a high degree of trait-and sex-specificity in chromatin architecture within the developing tissues (Fig. 3), indicating a mechanism by which co-option of shared regulators can rapidly evolve to scaffold distinct degrees of sexual dimorphism in a new body region.

Among the pleiotropic regulators identified here, we confirmed the role of the established master regulator *doublesex* in enabling coordinated, body-wide responses to the upstream cue of sex determination. Further, this study is the first to uncover trait- and sex-specific sets of Dsx binding sites in this species, opening the door for future mechanistic work to elucidate the trait-specific roles of this gene in horned beetles and other taxa (Fig. 4). Furthermore, the data generated here identified many new candidate regulators of sexually dimorphic development in this species that showed evidence of sex-responsive gene expression, putative binding to sets of sex-responsive CREs, or both. Through RNA interference, we functionally validated the role of one such transcription factor, *ventral veinless*, as a pleiotropic regulator of sex differences across multiple traits (Fig. 5). This finding highlights a non-canonical mode of action for a regulator of sexual dimorphism, as *vvl* exerts its role in the regulation of sex-biased development not via differential expression, but by utilizing binding sites that exhibit sex-responsive differential accessibility.

**Figure 5.**
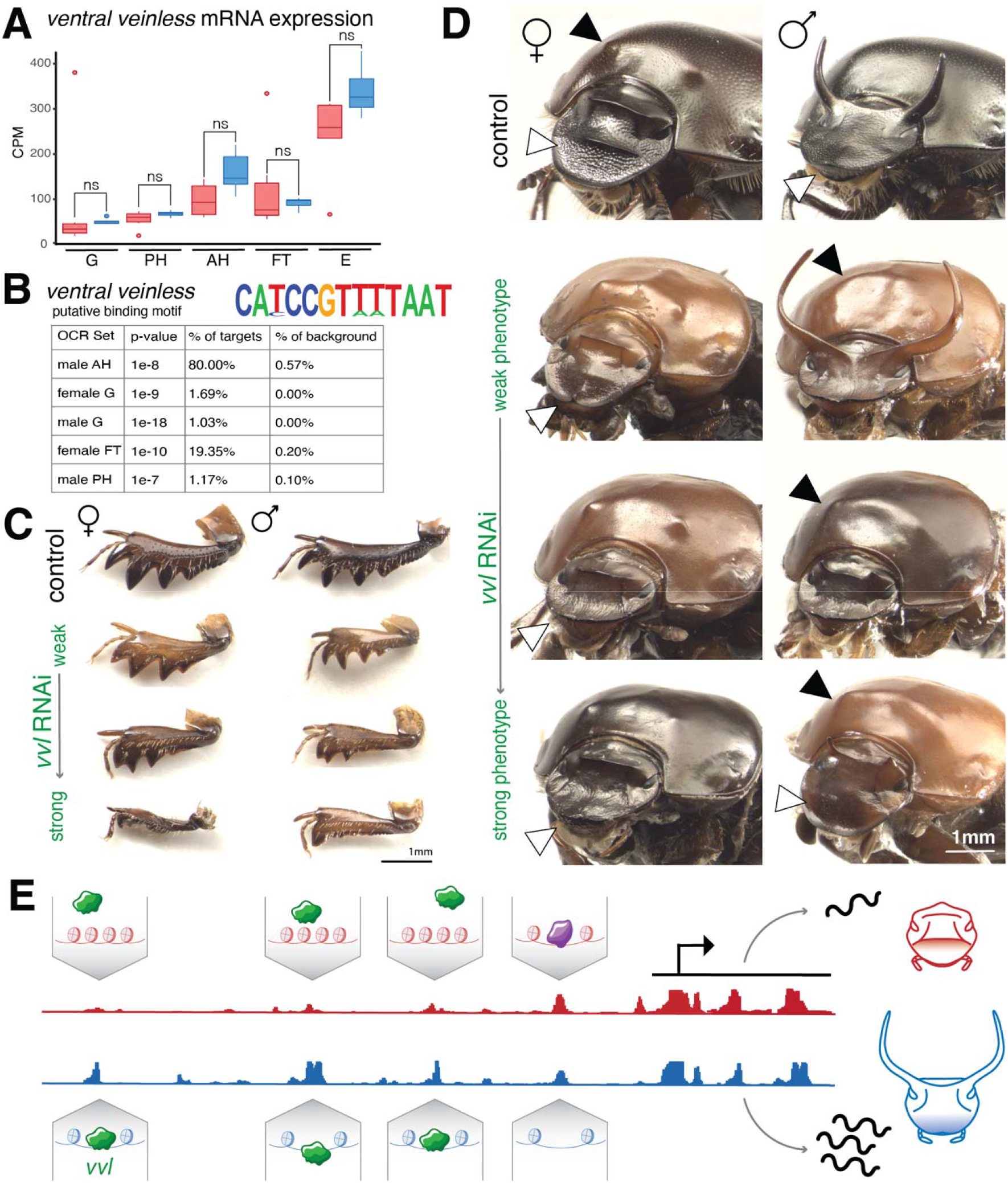
Transcription factor *ventral veinless* pleiotropically regulates sexually dimorphic trait development by binding unique sets of sex- and trait-responsive OCRs. (A) Putative binding motif of *Ot-vvl* is enriched in multiple sets of sex- and trait-responsive OCRs. (B) *Ot-vvl*^RNAi^ decreases fore tibia length and affects size and separation of tibial teeth in both sexes. (C) *Ot-vvl*^RNAi^ eliminates sex differences in the anterior head (white arrowheads) and prothorax (black arrowheads) but leaves the posterior head unaffected. (D) Model mechanism for a transcription factor that is *not* differentially expressed between the sexes (such as *Ot-vvl*, shown in green) regulating sexually dimorphic development exclusively through the differential accessibility of binding sites near its target genes, as opposed to transcription factors regulated via differential expression (shown in purple). G = genitalia; PH = posterior head; AH = anterior head; FT = fore tibia; E = elytra.

Together, this work furthers our understanding of the molecular mechanisms instructing the development of sex differences in somatic tissues, and provides some of the first data illustrating how transcriptional activity and chromatin remodeling interact to generate sexual dimorphism in a trait-specific manner. These findings contribute to a growing body of knowledge on how development integrates cues such as sex determination to enable highly similar genomes to yield different morphologies, allowing for adaptation to selective environments that affect each sex uniquely.

## Materials and Methods

### Beetle husbandry

*Onthophagus taurus* individuals were collected from Chapel Hill, North Carolina, USA and kept in laboratory conditions at 24^°^C as described by Shafiei et al. (2001). Six female and three male beetles were allowed to breed for one week, after which brood balls were collected and first instar larvae were moved to artificial brood balls in 12-well plates provisioned with grass-fed cow dung, following an established protocol (Shafiei et al 2001). Plates were checked every 12 hours for pupating individuals.

### RNA-seq Sample and Data Preparation

Upon the onset of pupation, live epithelial tissue was dissected from *O. taurus* pupae submerged in 0.05% Triton-X in phosphate-buffered saline (PBS) in sterilized glass dissection plates as described in Casasa et al. (2020). Posterior dorsal head tissue, anterior dorsal head tissue, the right elytron, the right foretibia, and the epithelial tissue comprising the two posterior-most ventral abdominal sclerites were all dissected from each male and female pupa and processed as separate samples. In total, we collected biological replicates of live epithelial tissues from 6 male and 6 female pupae for RNA-sequencing. Distinguishing the sexes of *O. taurus* pupae is straightforward, as the developing horns and genitalia of males become externally visible and recognizable at pupation. All dissected tissues were moved into ice-cold Trizol (Thermo Fisher Scientific) and stored at −80 °C until RNA extraction. Once all samples were collected, RNA extractions were performed using a Direct-zol RNA Miniprep kit (Zymo Research). Briefly, tissues were thawed to 4 °C, homogenized in the Trizol with RNase-free polypropylene pestles, and purified following the protocol on a Zymo spin column with a DNA digestion treatment. Total RNA quantity and quality was measured using an Agilent 2200 TapeStation system with an RNA ScreenTape Assay. A total of 60 (2 sexes x 5 tissues x 6 biological replicates) stranded RNA sequencing libraries were constructed with the TruSeq Stranded mRNA Sample Preparation Kit (Illumina) according to manufacturer instructions. Libraries were quantified using a Quant-iT DNA Assay Kit (Thermo Fisher Scientific), pooled in equal molar amounts and sequenced as paired-end reads using a 50-cycle P3 flow cell on the NextSeq2000 platform (Illumina).

### RNA-seq Data analysis

The resulting reads were cleaned using Trimmomatic v0.36 (Bolger et al 2014) to remove adapter sequences and perform quality trimming with the following parameters, 2:30:10 LEADING:10 TRAILING:10 SLIDINGWINDOW:4:15 MINLEN:20. The adapter-trimmed reads were used as input for Salmon (Patro et al 2017) to generate transcript abundance estimates indexed to the *O. taurus* v3.0 transcriptome (Davidson and Moczek 2024). Transcript TPM values from Salmon were imported into R and converted to gene-level counts using Tximport v1.4.0 (Soneson et al 2016). Statistical analyses of the RNA-seq data were carried out in R v.4.2.1. First, genes with extremely low read counts were removed (using a threshold of 3 counts-per-million required in at least 5 biological replicates). Comparison of biological replicates was performed by calculating a matrix of pairwise Euclidean distances between samples. This method identified one female elytra sample as a distinct outlier not only from its tissue type, but from all other samples in the analysis (supplementary figure 5), and was therefore removed from the sample set.

To explore expression differences between sexes and tissues, a DESeq2 dataset was generated using a multifactor design that incorporated “sex” and “tissue” as factors using the *group* function (Love et al 2014). After fitting this model to the data, differential expression comparisons were made using the DESeq2 *contrast* argument to specify pairwise comparisons of sample groups. For each tissue type, “female” was set as the reference and “male” was set as the comparison sample type, and differential expression was assessed in this pairwise manner for all five traits. Transcripts with a Benjamini and Hochberg false discovery rate <0.1 and log2 fold-change of <0 or >0 were considered to be differentially expressed for a given contrast, in an effort to be more exploratory than stringent in assessing sex-responsiveness in the transcriptome. Venn diagrams of sex-responsive gene sets were generated using InteractiVenn (Heberle et al 2015). The resulting sets of statistically significant sex-biased genes were used for downstream gene ontology term enrichment analyses. To assign GO terms, BLAST (Altschul et al 1990) was used to query all genes in the *O. taurus* transcriptome against the UniProtKB/SwissProt database of reviewed eukaryotic gene annotations (UniProt Consortium 2025). Enrichment analysis was performed using *topGO* in R (Alexa and Rahnenfuhrer 2024). Reported p-values are not corrected for multiple testing, as the terms within the GO database, and thus the statistical tests are not fully independent from one another (Li et al. 2021).

### ATAC-seq Sample and Data Preparation

We collected biological replicates of live epithelial tissues from 5 males and 5 females for ATAC-sequencing. Upon the onset of pupation, live epithelial tissue was dissected from O. taurus individuals in molecular-grade 1X PBS in sterilized glass dissection plates. Again, posterior dorsal head tissue, anterior dorsal head tissue, the right elytron, the right foretibia, and the epithelial tissue comprising the two posterior-most ventral abdominal sclerites were all dissected from each male and female pupa and processed as separate samples. After dissection, 50,000 cells (determined by counting DAPI stained cells in a hemocytometer) were immediately transferred to a 1.5 mL conical tube and processed for ATAC-sequencing (Buenrostro et al 2013) using the Omni-Seq protocol (Corces et al 2017). Sequencing libraries were generated by amplifying open chromatin fragments with PCR, after having determined the optimal number of amplification cycles with qPCR, as described by Buenrostro et al (2015). Samples were sequenced on an NextSeq550 platform (Illumina) with a 75-cycle high output kit to a minimum depth of 10.7 million paired-end reads per sample, averaging 25.4 million reads per sample. ATAC-seq libraries were trimmed and filtered with *Trimmomatic* v. 0.39 (parameters: leading:10 trailing:10 slidingwindow:4:15 minlen:25) (Bolger et al 2014). Filtered, paired-end ATAC-seq reads were then aligned to the *O. taurus* genome assembly (Davidson and Moczek 2024) with *Bowtie* v.2.2.5 (Langmead and Salzburg 2012). Alignments were filtered for minimum mapping quality (MAPQ) score of 20 using *samtools* (cite) and PCR duplicates were removed with the *filterdup* tool from macs2 v.2.2.6 (Zhang et al 2008). ATAC-seq peaks (hereafter open chromatin regions: OCRs) were called with *macs2* using the parameters:—nomodel—keep-dup = auto —shift 100—extsize 200 -g 2.9e8 -f BAMPE -q 0.05 (Zhang et al 2008). All OCR sets across sample types were merged into a consensus set of 175,884 OCRs with the *bedtools* v.2.30.0 merge command, and peak accessibility counts were calculated with the *bedtools multicov* command (Quinlan and Hall 2010).

### ATAC-seq Data analysis

Statistical analyses of the ATAC-seq data were carried out in R v.4.2.1. Peaks with read counts below 3 counts-per-million required in at least 5 biological replicates were removed, leaving a final set of 88405 high-quality OCRs for downstream analysis (S2 Data). To account for X chromosome copy number differences between males and females as is becoming standard in the field (Pal et al. 2019, Su et al 2025), read counts mapping to the X chromosome in males were doubled. Differential accessibility analyses between males and females for each body region were calculated in DESeq2 v.1.36.0 (Love et al 2014), with female sample groups designated as the reference, and male groups as the comparison, where significantly differentially accessible OCRs were called as having false-discovery rate support <0.1 and and log2 fold-change of <0 or >0. Separate DESeq2 analyses were performed on the full dataset to assess pairwise differential accessibility between traits.

Sets of statistically significant sex-biased OCRs within each trait were used for downstream analysis. Transcription factor binding motif prediction and motif enrichment analyses were carried out in homer v.4.11 (Heinz et al 2010) for each OCR set of interest on a background set of all pupal epithelial OCRs identified in this study using the *findMotifsGenome*.*pl* script with the parameters: -50,50 -mset insects -fdr 10. Putative transcription factor identities were assigned using the *compareMotifs*.*pl* script to search for similar motifs across the program’s database of validated insect motifs (derived primarily from work in *Drosophila*).

### Double-stranded RNA synthesis, injection, and phenotype assessment for RNA interference

A target region of the gene *Ot-ventral veinless* was chosen using a BLAST algorithm to query 250bp regions of the gene against the O. taurus transcriptome and selecting one with zero off-target hits (Supplementary Table 1). An oligonucleotide construct and T7 RNA polymerase primers were designed using the reference genome for the species and ordered from Integrated DNA Technologies, Inc. Synthesis of double-stranded RNA (dsRNA) for gene knockdown via RNA interference (RNAi) was performed using a protocol optimized for coleopteran larvae (Philip & Tomoyasu 2011). Briefly, PCR was performed to anneal T7 RNA polymerase binding sequences to the ends of the gene fragment construct to generate a DNA template, which was purified using the Qiagen QIAquick PCR Purification kit. In vitro transcription was performed using an Ambion MEGAscript T7 kit to generate dsRNA from the template, which was purified using the Ambion MEGAclear kit with an ethanol precipitation step. For injection, the dsRNA construct was diluted to the target concentration (between 1-0.005 µg/µL, see Supplementary Table 1) with injection buffer (recipe from Philip & Tomoyasu 2011). A Hamilton brand syringe and small gauge removable needle (32 gauge) were used to inject a 3µl dose of dsRNA through the abdominal cuticle into the hemolymph of each third-instar larva to initiate whole-body RNAi. Control individuals were randomly selected from each round of developing larvae and injected with pure injection buffer.

Previous work in dung beetles has shown that injection of pure buffer alone can serve as a suitable control for dsRNA injection, as neither buffer-injected nor nonsense RNA-injected adults have been documented to show any detectable phenotypic differences compared to wildtype adults (Moczek & Rose 2009, Simonnet & Moczek 2011, Linz et al. 2019). After eclosion, adult animals were sacrificed and stored in 70% ethanol before analyzing and comparing the phenotypes of RNAi-injected and control-injected beetles. Representative individuals from each treatment group and sex were photographed using a Leica MZ16 microscope with a PLANAPO 2.0x objective and a Leica S8APO microscope and a PixeLINK PL-D7912CU-T camera; multiple photos of each sample were taken across different planes of focus and merged using Adobe Photoshop.

## Acknowledgements

The authors would like to thank Dr. David Merritt and the staff at the Indiana University Center for Genomics and Bioinformatics for assistance with sequencing platforms, Dr. Phil Davidson for advice on sequencing library preparations and computing clusters, Yongsoo Choi and Miranda Towse for help monitoring many plates of larvae, and the IU Eco-Evo-Devo group for feedback on earlier drafts of the manuscript. This work was supported by generous funding from the National Science Foundation [Grant no. 2243725 and 1901680 to APM] and was performed while EMN was funded by the National Institutes of Health [T32-HD049336].

## Data Availability

Raw sequencing data are available through the NCBI SRA (RNA-seq: BioProject number PRJNA1425507 https://www.ncbi.nlm.nih.gov/bioproject/PRJNA1425507, ATAC-seq: PRJNA1425599 http://ncbi.nlm.nih.gov/bioproject/PRJNA1425599). Scripts for read processing and mapping, R code, and data files needed to replicate all analyses are available on Github: https://github.com/ericanadolski/Otaurus_sexual_dimorphism

