## Supplementary-figures-tables for "Trait-specific chromatin architectures channel pleiotropic genes toward sexually dimorphic development in horned beetles"

### Supplementary Data

Supplementary table 1. Differential gene expression data – separate attachment

Supplementary table 2. Differential chromatin accessibility – sex-responsive OCRs – separate attachment

Supplementary table 3. GO term enrichment data – separate attachment

Supplementary table 4. Motif enrichment data – separate attachment

Supplementary table 5. Differential chromatin accessibility – trait-responsive OCRs – separate attachment

Supplementary table 6. Trait- and sex-responsive OCRs within 20kb of *doublesex*

Supplementary Table 7. *Otau-ventral veinless* BLAST identification and RNA interference target region

| **BLASTp Query** | **BLASTp Hit** | **e-value** | **bit score** | **Hit Gene Sequence (RNAi target region underlined)** |
| --- | --- | --- | --- | --- |
| *D. melanogaster ventral veinless* | jg3443.t1 | 2e-64 | 215 | ATGGCGGCAGCAACATACCTGCCGACGAGTAGTGCCCTGACATCAGATGTGGACGGTGGAGTAGTGGTCGGCATGAACGGGATGAACATCGCTGGAGGTTATCATAGTAGTAACTCGCCCAGATCGGGCGTTGATAGTGACATGAAGTACCTTCCGCATCATCAACACCATCATCATCACCAGCATCATATGCCCGCGTCGCCGTCGCCTTCCGGGCCAGCAATGATGAATCCTTGGGTTAGTTTACAATCGTCCGATCCGTGGAATATGCCAATTCACCATCATCATCAGGATATAAAACCTTTGGCGCAACACGCCGAAATCTTACACAGACAGCAACAGCAACAACTTAGTTCACCGCATCACCATGGTTGGTCGATGTCACAACCTTCTCATTATAATCCCGGCGCTGGTAGCCCATTGCAAACCGTTAACGGTATGCTATCTCAGTACCCACCGACGCCGCAACTTCATCATTCGTTACACCGCGAACTCCAAAGTCCACACCACCCGCAGCATCCCGGAGATCGAGATAGCGTCGGTGAAGATGAAACCCCGACAAGCGACGATCTAGAAGCCTTCGCCAAACAATTTAAACAAAGACGAATAAAGCTTGGTTTCACCCAAGCTGATGTTGGTTTAGCATTGGGTACTCTTTACGGAAACGTTTTCTCTCAAACAACAATTTGTCGATTTGAGGCTCTCCAATTGAGTTTCAAAAATATGTGCAAACTTAAACCTTTATTACAAAAATGGCTCGAAGAAGCCGATTCGACGACCGGTTCCCCAACATCGATCGACAAAATAGCGGCGCAGGGTCGAAAGCGGAAAAAGCGCACCAGTATCGAGGTATCTGTAAAAGGGGCCCTAGAACAACATTTTCATAAACAACCGAAACCTTCAGCACAAGAAATTAGCAGCTTAGCGGATAGCCTTCAATTAGAAAAGGAAGTTGTAAGAGTTTGGTTTTGCAACCGACGGCAAAAAGAGAAAAGGATGACTCCGCCGAATACTCTCGGTCCAGATATGATAGACAACCTTCCACCGAGCGGTCATATCCACCCAAGTTACGGCCATCACCCAACGGATATGCACGGATCACCGATGGGTGGACAACATTCCGTTTCCCACAGCCCCCCAATGTTATCCCCGCAAGGGATGGGACACCAGCTTACGGCCCACTAG |

Supplementary Table 8. RNA interference sample sizes and penetrance.

| Target Gene | Dose (ug/ul) | # Treated Larvae | # Dead | # Surviving Adults | # Adult Males | Penetrance of Male Phenotype | # Adult Females | Penetrance of Female Phenotype |
| --- | --- | --- | --- | --- | --- | --- | --- | --- |
| buffer | n/a | 34 | 6 | 28 | 10 | n/a | 18 | n/a |
| *Ot-vvl* | 1.0 | 20 | 17 | 3 | 2 | 100% | 1 | 100% |
| *Ot-vvl* | 0.5 | 23 | 22 | 1 | 1 | 0% | 0 | n/a |
| *Ot-vvl* | 0.25 | 40 | 36 | 4 | 4 | 50% | 0 | n/a |
| *Ot-vvl* | 0.025 | 124 | 97 | 27 | 15 | 53.3% | 12 | 75% |
| *Ot-vvl* | 0.01 | 54 | 45 | 9 | 7 | 28.5% | 2 | 50% |
| *Ot-vvl* | 0.005 | 90 | 38 | 52 | 21 | 42.8% | 31 | 45.2% |

Supplementary Figure 1. Principal component analysis plots performed on full sample sets, with samples plotted along PC1 (x axes) and PC2 (y axes) for the RNA-seq experiment (left) and ATAC-seq experiment (right).

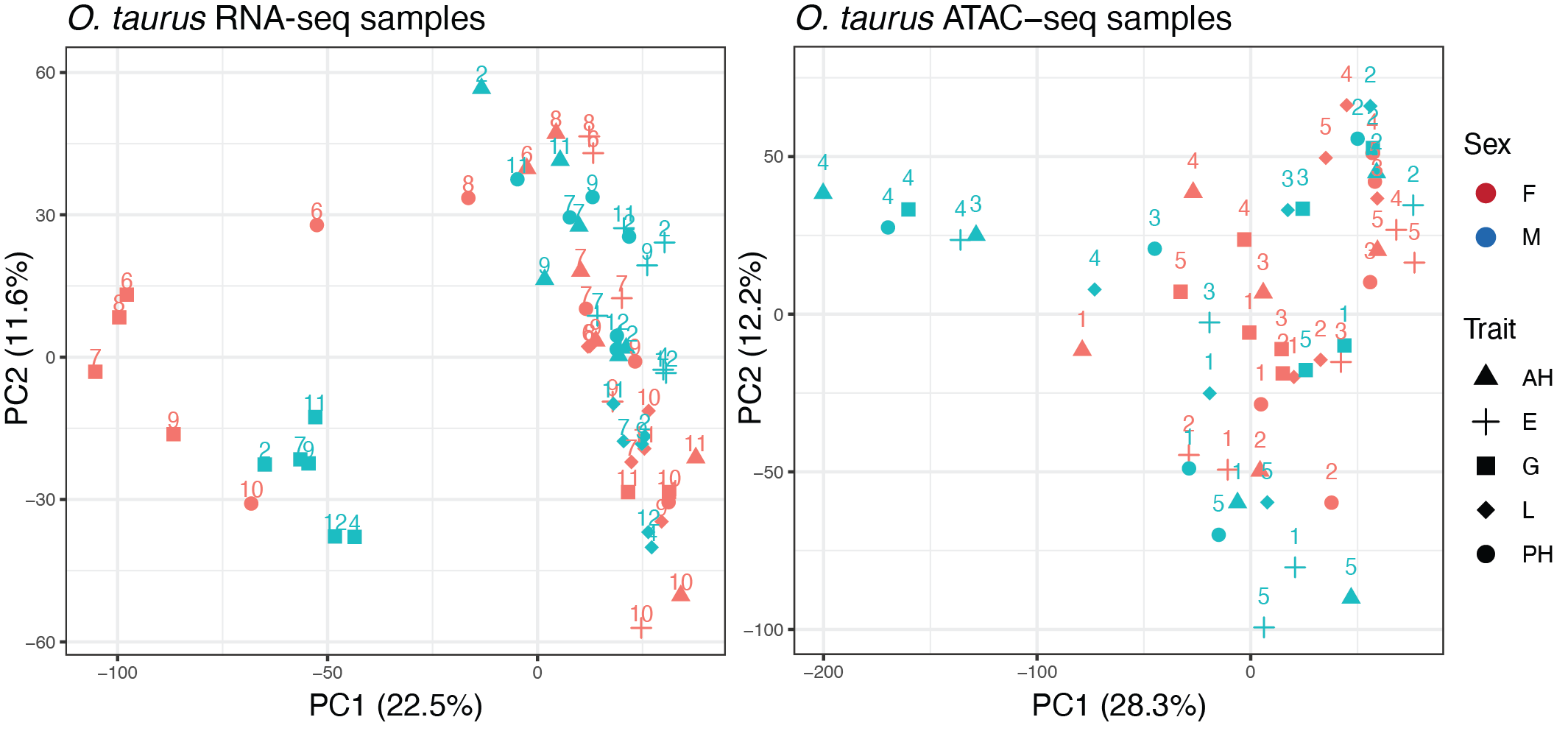

Supplementary Figure 2. Heatmaps showing hierarchical clustering of samples and gene expression levels for the differentially expressed genes called by DESeq2 for each trait.

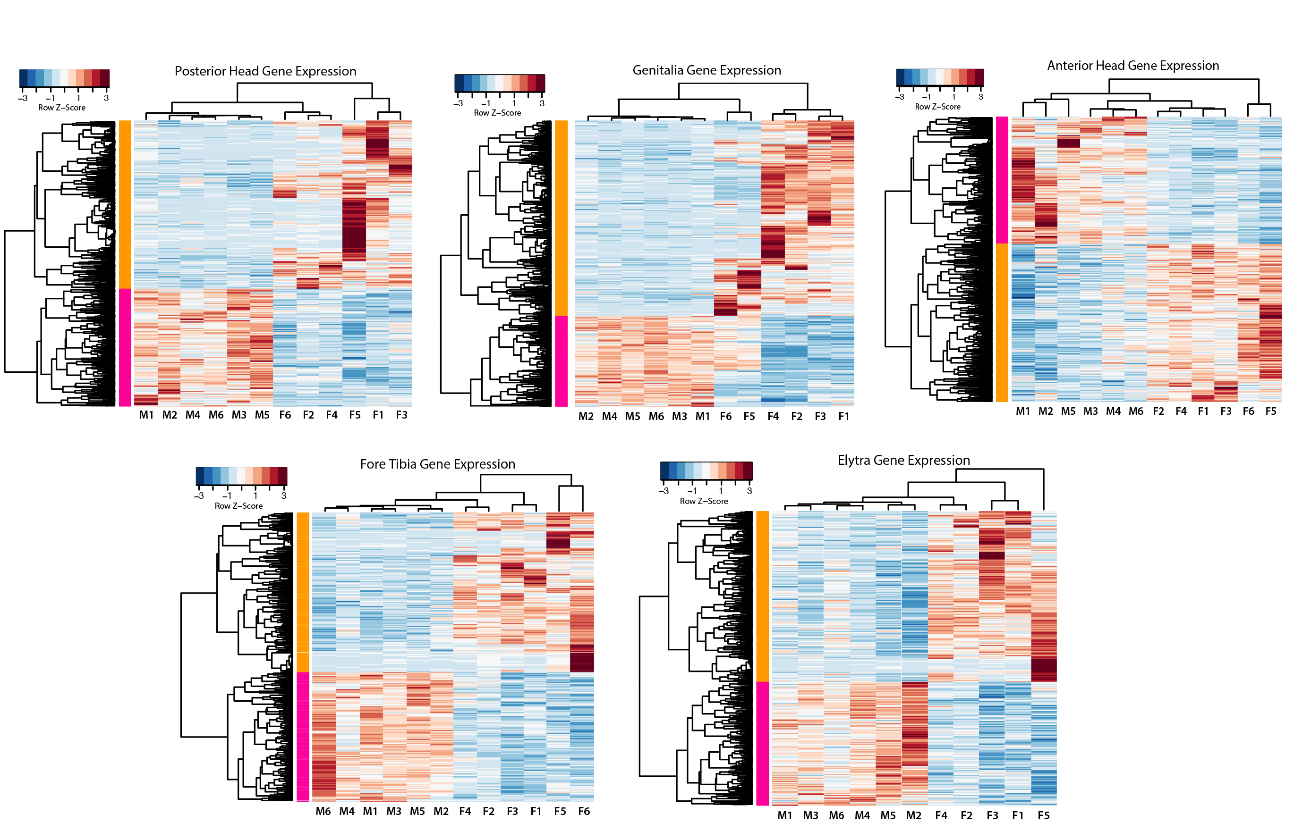

Supplementary Figure 3. Heatmaps showing hierarchical clustering of samples and open chromatin region (OCR) peak read counts for the differentially expressed OCRs called by DESeq2 for each trait.

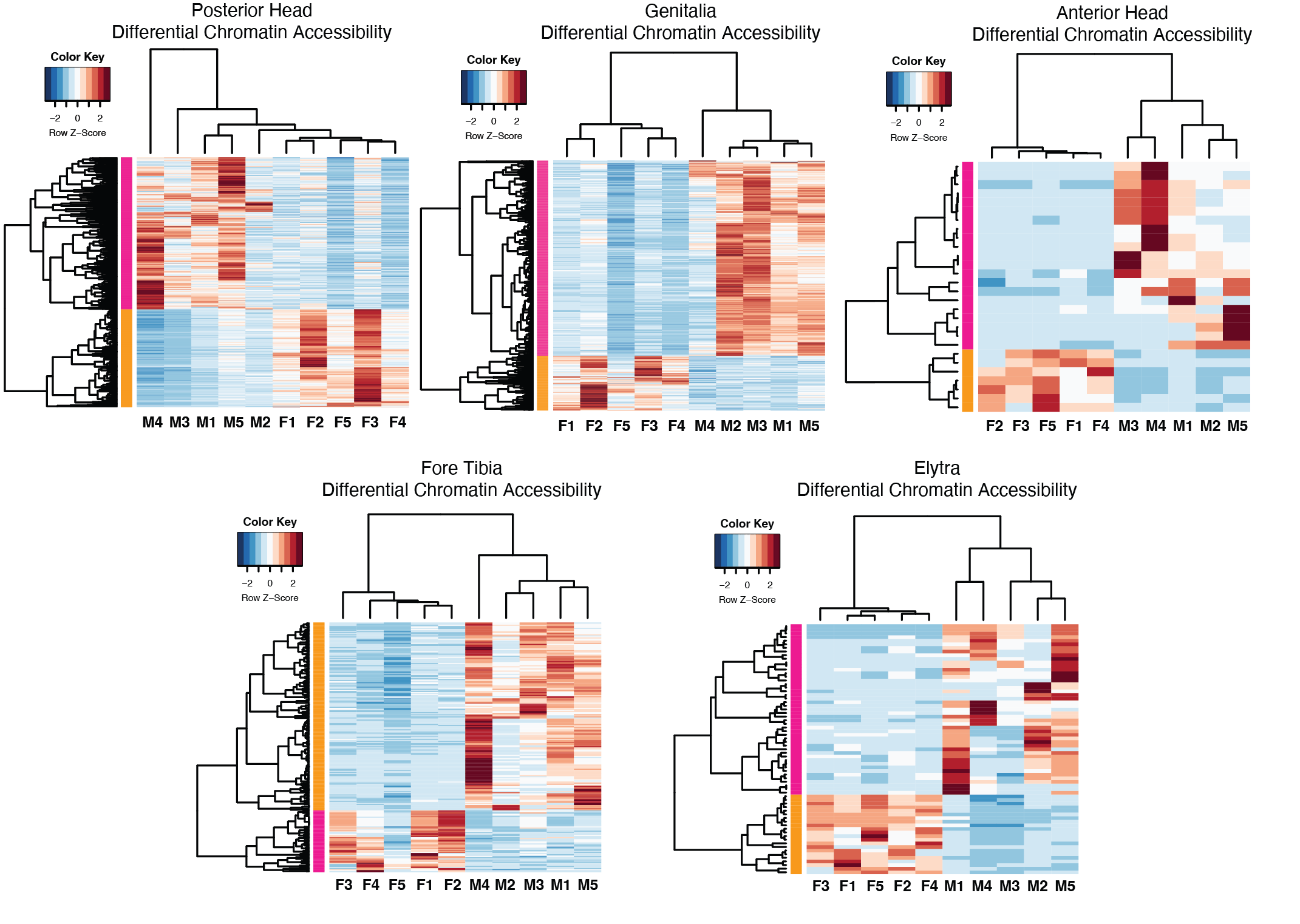

Supplementary Figure 4. Additional phenotypes of *Otau-vvl* RNAi. (A) Dorsal head images of control-injected and *Otau-vvl* RNAi beetles highlighting the differences in fine-scale cuticle morphology. (B) T3 wing of a control-injected (top) and *Otau-vvl* RNAi beetle (bottom) dissected away from the body, highlighting defects in wing venation and a decrease in overall wing length. (C) Full-body view of control-injected (left) and *Otau-vvl* RNAi beetle (right) highlighting defects in T2 elytra and T3 wing sclerotization and positioning (white arrowheads).

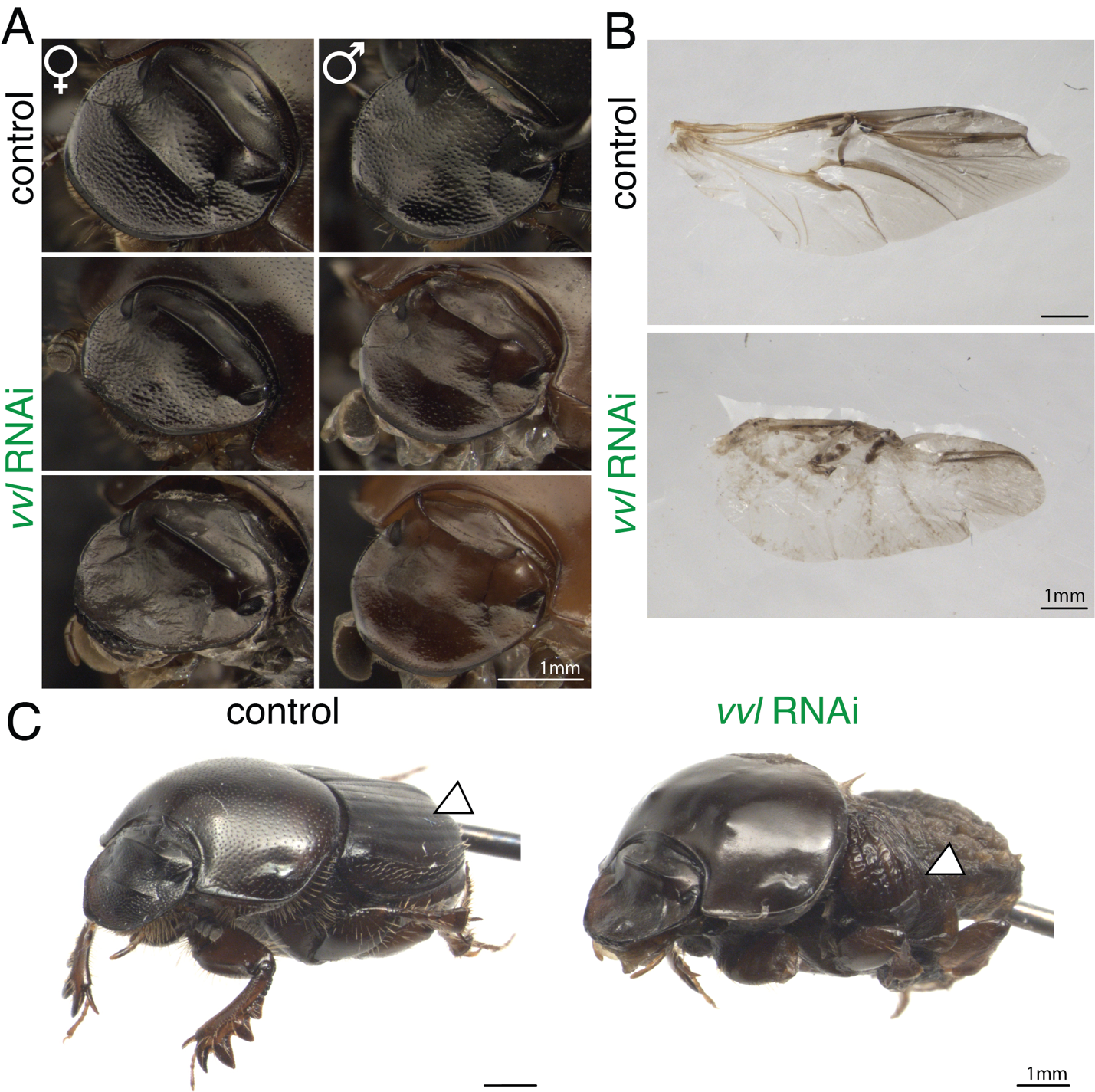

Supplementary Figure 5. Analysis of RNA-seq outlier sample. (A) Hierarchical clustering heatmap of elytra gene expression samples and differentially expressed genes called by DESeq2 showing Ot-F11-E as an outlier to all other female and male elytra biological replicates. (B) Bar chart tallying the number of genes called as differentially expressed with the outlier sample included (white bars) and excluded (grey bars).

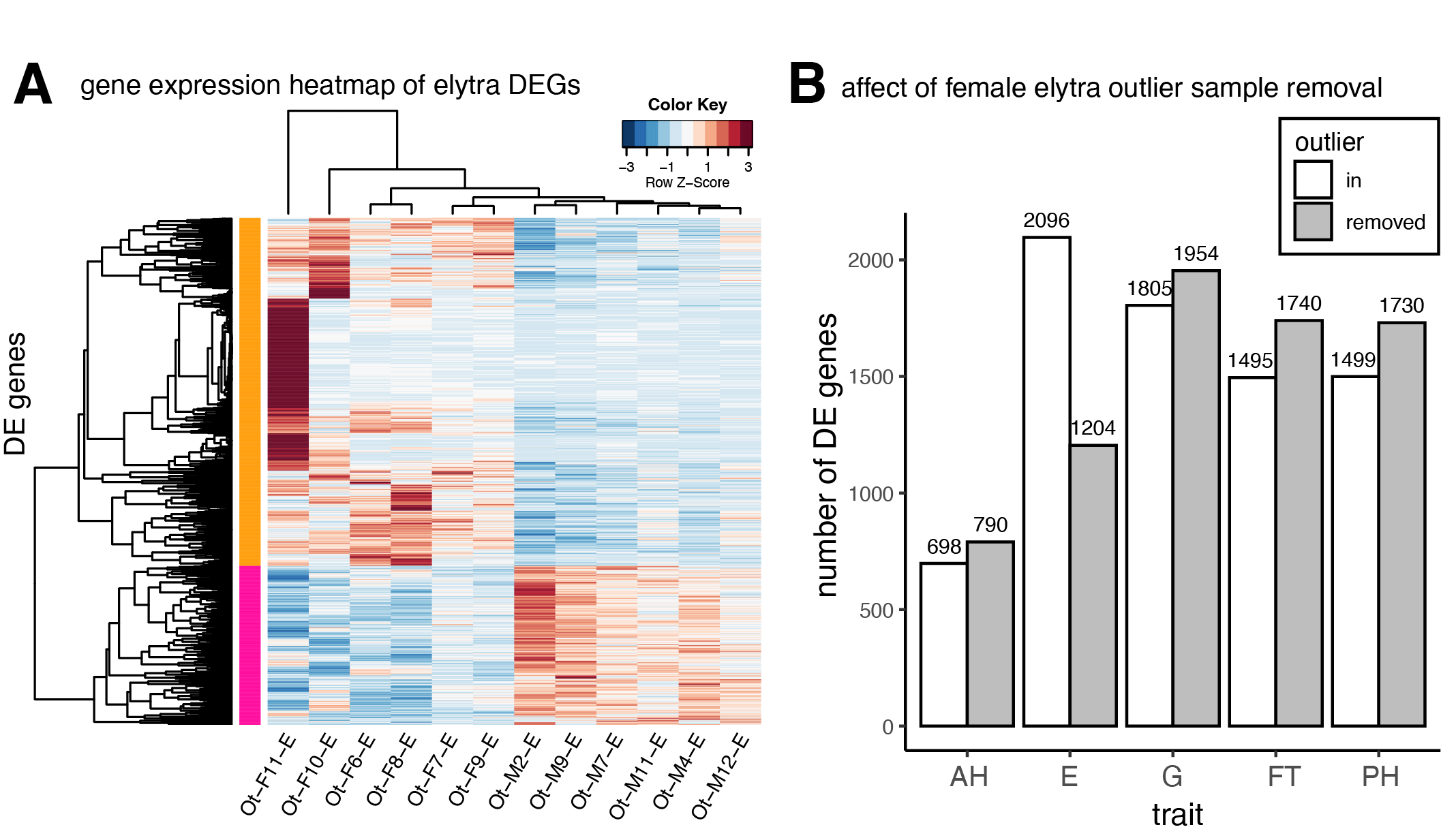
